# Data-driven simple agent-based model of scratch assays on healthy and keloid fibroblasts

**DOI:** 10.1101/2024.04.02.587674

**Authors:** Stéphane Urcun, Gwenaël Rolin, Raluca Eftimie, Alexei Lozinski, Stéphane P.A. Bordas

## Abstract

In this study we propose a novel agent-based model to reproduce and propose new hypotheses on the biological mechanisms of cell-cell interactions and cell migration from data obtained during scratch assay with healthy and keloid fibroblasts. The advantage of the agent-based model we propose in this paper lies in its simplicity: only three governing parameters. We conducted a parametric sensitivity analysis and we incorporated the evaluation of contact inhibition of locomotion, aligning with the observed loss during malignant invasion. To study invasion modalities, we conducted *in vitro* wound healing assays using healthy and pseudo-tumoral (keloid) fibroblasts under diverse conditions: control, macrophage type 1 secretome, and macrophage type 2 secretome. Mitomycin inhibition of proliferation isolated the contribution of migration to wound filling. Our agent-based mathematical model describes configurations based on our microscopy imaging and statistical data, which enables quantitative comparisons between our experimental and numerical results. Calibration and evaluation were performed on the same experiments, enriched by external datasets. With only three governing parameters, our model not only demonstrated good agreement (8.78% to 18.75% error) with external evaluation datasets for all experimental configurations but also provided us with a nuanced understanding of keloid fibroblast behavior during wound healing, especially regarding contact inhibition dynamics.

## Introduction

Skin wound healing is a physiological and well-orchestrated process of tissue repair. In some pathological situation, this process is dysregulated and leads to an over-accumulation of fibrotic tissue. Keloid is an example of such pathology and is characterized by a tissue invasion outside the wound border [1]. Keloid fibroblast is one of the main cell actor engaged during keloid development. All along the process, fibroblasts are also in close interaction with immune cells (*i.e* macrophages) that regulate their behavior through secreted biological factors. Compared to normal fibroblasts, keloid fibroblasts demonstrate an exacerbate capacity of extra-cellular matrix synthesis, proliferation and migration [2–5]. To study collective and individual cell’s migration involved in wound healing, *in vitro* scratch assays are widely used [6, 7]. The reader may refer to fibroblasts and keratinocytes co-cultured in enhanced serum in Walter *et al*. [6], or dermal fibroblast migration through microfluidic design and pressure induced wound in Shabestani *et al*. [8], as a few examples of this rich experimental designs where cells or biological factors can impact on fibroblasts migration. In our study, the presented scratch assays only focus on one cell type with two modalities: healthy or pathological (keloid) fibroblasts. The interaction and migration of both type of cells were followed in classical culture condition or in the presence of polarized macrophages secretome to reproduce a pro-inflammatory (M1 polarization) or a pro-fibrotic (M2 polarization) micro-environment, as previously described by Setten *et al*. [9] and Dirand *et al*. [10]. Our sober design aims to benchmark healthy dermal fibroblast behavior and compares it to keloid fibroblasts. Importantly, this experimental design is well-suited for mathematical modeling. By reducing the complexity, these assays allow likewise to reduce the complexity of the mathematical model, therefore reducing the computational time of its execution and calibration.

Agent-based modeling is one mathematical method to approach cellular behavior, considering each cell individually, interacting with others. Their characteristics and the rules of interaction constitute the agent-based model. This approach allows a large freedom of modeling, that we can divide on categories: free or lattice-based simulation, with or without description of subcellular compartments, and hybrid, *i.e*. that admit continuous fields - such as chemical signaling - within the agent-based model. In this study, we choose a specific lattice-based method, the Cellular Potts Model (CPM), developed by Graner and Glazier [11]. If a regular grid defines the computational domain, an agent is not confined in one element: the geometry of each agent is defined by a set of elements, allowing for its detail description. Therefore, photographs of *in vitro* experiments may be translated, pixel-wise, in the CPM computational domain. This opens the way to a quantitative comparison between the surface occupied by the fibroblasts in the experiments and in the numerical results. The major computational framework used for CPM is CompuCell 3D (CC3D) [12], sustained by a large community of developers.

Despite the richness of the agent-based modeling field, the computational literature which focuses on fibroblast is quite sparse. It mainly includes interaction with other cellular population, such as the study of Andasari *et al*. [13], an hybrid model where keratinocytes migration is described by the means of endothelial growth factor secreted by the fibroblast population. However, this rich model admits 27 parameters, which lead to a heavy calibration process and may provoke difficulties for a numerical analysis. Closer to our study, the early works of Dallon and Sherratt [14] in 1998 studies influence of collagen orientation on fibroblasts migration, and in 2000, Cobbold and Sherratt [15] designed their keloid mathematical model on inflammation and cell densities. An other example of fibroblast’s behaviour studied by mathematical modeling is the coupled *in vitro, in vivo, in silico* study on a large experimental dataset of Rognoni *et al*. [16], on the interaction between fibroblasts, that proliferate or migrate, and ECM architecture and deposition. Additionally, the cancer modeling community studied the role of cancer-associated fibroblasts under the perspective of wound healing, see the work of Norton *et al*. [17], and with an agent-based approach in Heidary *et al*. [18].

The quantification of experimental data is a crucial challenge for robust simulation. As claimed in a later work of J. Uitto and M. Tirgan [19]: “it appears that the biggest obstacle to development of optimal treatment of patients with keloids is the lack of data-driven treatment pathways”. The two studies of Johnston *et al*. [20, 21] are clear examples of its advantages for comparison and interpretation between experimental and numerical results. In this study, we focus on fibroblast-fibroblast interaction during their migration in scratch assays. According to the quantification of experimental data, the fibroblast characteristics - the healthy ones and their pathological deviations – are directly extract from the experimental data, such as their principal length *(i.e*., the greater segment circumcised in the fibroblast body crossing its nucleus) or their surface on the culture plate. In the agent-based model, fibroblast to fibroblast interaction is mainly govern by their surfaces in contact. This phenomenon was first unveiled by Abercombie *et al*. [22] in 1953 and coined as contact inhibition of locomotion (CIL). The contact micro-structures of fibroblast are in-depth studied by Chen *et al*. in [23]. The study of CIL is pursued by Mayor *et al*. in [24], they highlight that pathological invasive behavior which occurs in cancer or keloid is characterized by the loss of CIL. This concept belongs to collective cells migration modeling, reviewed in [25].

This study aims to a step-wise quantitative approach of agent-based modeling of fibrosis: its applies first to healthy human dermal fibroblasts, then to keloid fibroblasts. This step-wise approach will lead to a robust in silico modeling that authorizes to simulate multiple experimental scenarii of fibrosis onset, without expense of biological material. Importantly, the quantification of the experimental data allows for i) informing as much as possible the input parameters of the mathematical model, such as relevant initial guesses, ii) reducing its complexity and the computational cost of its calibration, iii) rigorously computing the error between the mathematical model and the experiments.

We first present the experimental data and the process of their quantification. Then, we present our CPM. Through variance-based sensitivity analysis, we reduce the model calibration to governing parameters only. The experimental dataset is divided into two parts, the first half is used to calibrate the governing parameters, the second half is used to evaluate the model calibration.

## Material and Model

### Experimental data description

#### Secretome preparation

Macrophages secretome were produced via the following process. First, human monocytes were isolated from leukocyte-platelet concentrates. On one hand, M1 macrophages were generated from monocyte by treating them during 7 days with GM-CSF (50 ng*/*mL) and IFN*γ* (50 ng*/*mL), followed by a 24h treatment with 100 ng*/*mL LPS. On the other hand, M2-like macrophages were generated from monocyte by treating them during 7 days with M-CSF and IL-4 at 50 ng*/*mL, followed by a 24h treatment with IL-4 (20 ng*/*mL) and IL-13 (20 ng*/*mL). After polarization, both M(LPS) and M(IL-4, IL-13) were rinsed with buffer and cultured in fresh medium during 24h to produce M1 and M2 secretome (*i.e*. medium containing molecules secreted by M1 or M2).

#### Biological assay

For scratch wound assay, we used either healthy or keloid dermal fibroblasts extracted from human sample. Fibroblasts were seeded at 15K cells/wells in ImageLock 96-wells plate (Sartorius). After 24h adhesion, cell proliferation was blocked using mitomycin C (5*μg/*mL) for extra 24h. This step is crucial to be sure that wound filling is only due to cell migration and not a combination with cell proliferation. Cell monolayers were then wounded using WoundMakerTM device (Sartorius). After wounding, cells were rinsed twice with buffer before cell treatment with normal culture medium (Ctrl) or secretome generated from type-1 and type-2 macrophages (M1 and M2). Wound closure was followed over time with real-time microscope (IncuCyte S3TM, Sartorius).

#### Ethical statement

Keloid tissues were obtained from patients undergoing reductive plastic surgery performed at Maxillo-Facial Surgery Department of the University Hospital of Besançon (CHU of Besançon, France). All included patient provided informed consent and the study was conducted in accordance with the ethical standards, namely the Declaration of Helsinky. This work was ethically approved by the French Regulatory Agency (ANSM), ethic committee (CPP Sud-Ouest and Outre-Mer I), and was registered on clinicaltrial.gov as “SCAR WARS” (NCT03312166).

Leukocyte-platelet concentrates were obtained from healthy volunteers at the French Blood Establishment (EFS Bourgogne Franche-Comté, Besançon, France) after informed consent signature (authorization number #AC-2015-2408).

### Experimental data quantification

Data used in this study are constituted of 2D culture plates photographs of 1.46 *×* 1.97 mm size, of 1176 *×* 1584 resolution, which gives a pixel length of 1.24 *μ*m. For the sake of simplicity, all the lengths and surfaces are given in pixel (px) and pixel square (px^2^). The pictures are taken at *T*_0_, *T*_0_ + 20 H and *T*_0_ + 36 H. Experiments are divided in 3 types, Control, M1 secretome and M2 secretome. In this study, we reproduce seven assays, one with healthy fibroblasts HF-Ctrl, and six with keloids fibroblasts, namely CtrlA, CtrlB, M1A, M1B, M2A and M2B.

As far as we know, there is not automatic counting and segmentation adapted to the specific category of fibrotic cells. Although automatic counting and segmentation of cell is a well-studied field, it is applied with restricted conditions. Efficient algorithms are applied on histopathological staining [26, 27], but they cannot be used with grey level microscopy images. Other algorithms succeed in automatic counting and segmentation with microscopy images [28], but only for ameboid cells or cells with well-defined shapes and without overlapping. Therefore, all the data presented in this study were manually extracted. Our group is currently developing an algorithm specific to fibroblast population. The data presented in this study constitutes a part of the labeled data used for the development of this algorithm.

For each of the 8 experiments, the following treatment is applied with ImageJ [29]:

- At *T*_0_, all the healthy or keloid fibroblasts’ nuclei are labeled and their coordinates exported. All the labeled fibroblasts are contoured with the polygon tool. See Fig.1 A, B, C.
- We define, in the experimental images, a domain between the lines define by (*x* ∈ [0; 1176] px, *y* = 600px) and (*x* ∈ [0; 1176] px, *y* = 1000px) (see red lines Fig.2B,C). This domain, always empty at *T*_0_, corresponds to the wound domain to close by the fibroblasts. The fibroblasts inside this domain at *T*_0_ + 20 H and *T*_0_ + 36 H are labeled, contoured, and their surfaces measured (see the process Fig.2A, B, C). This provides the percentage of the wound closure displayed Table 1.

**Fig 1.**
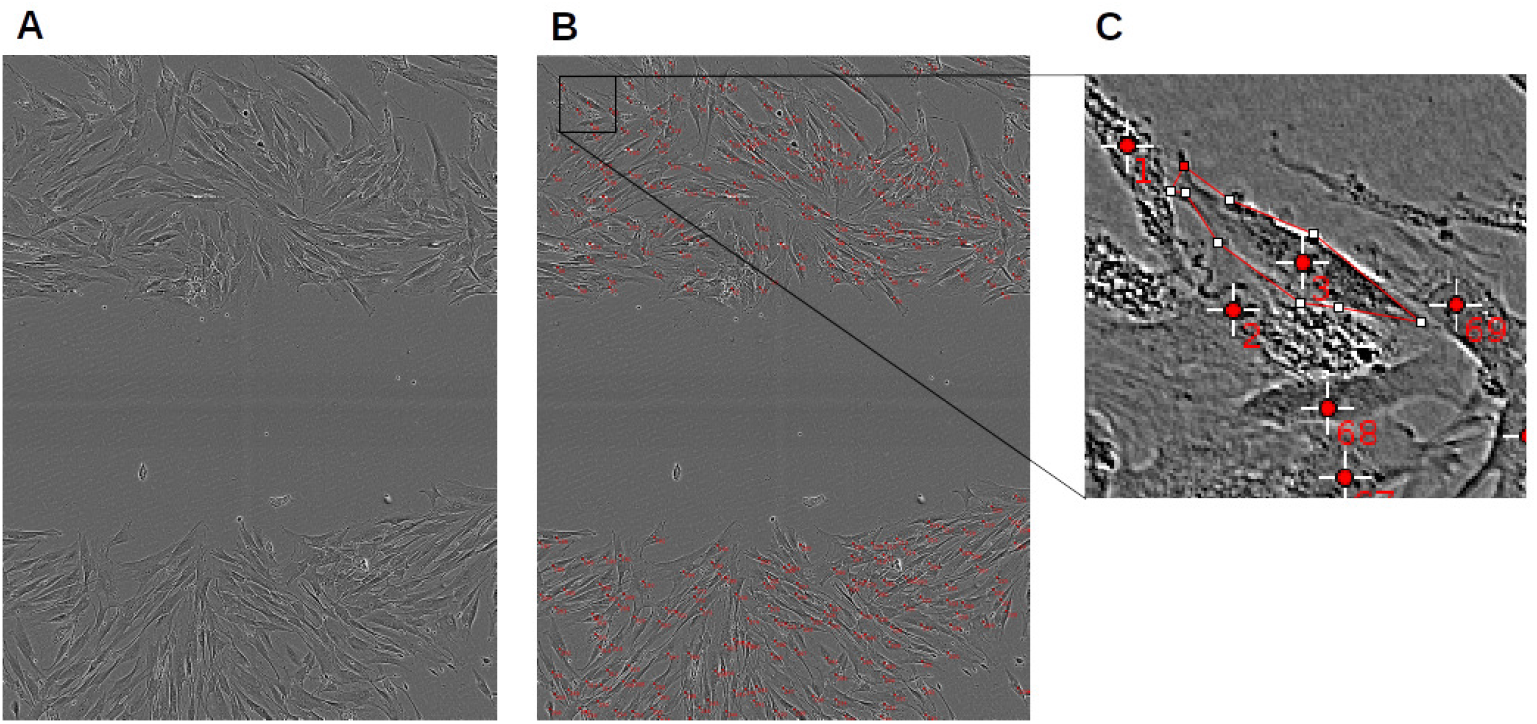
Experimental data quantification: CtrlA initial conditions. At *T*_0_, all the healthy or keloid fibroblasts’ nuclei are labeled and their coordinates exported. All the labeled fibroblasts are contoured with the polygon tool of ImageJ [29].

**Fig 2.**
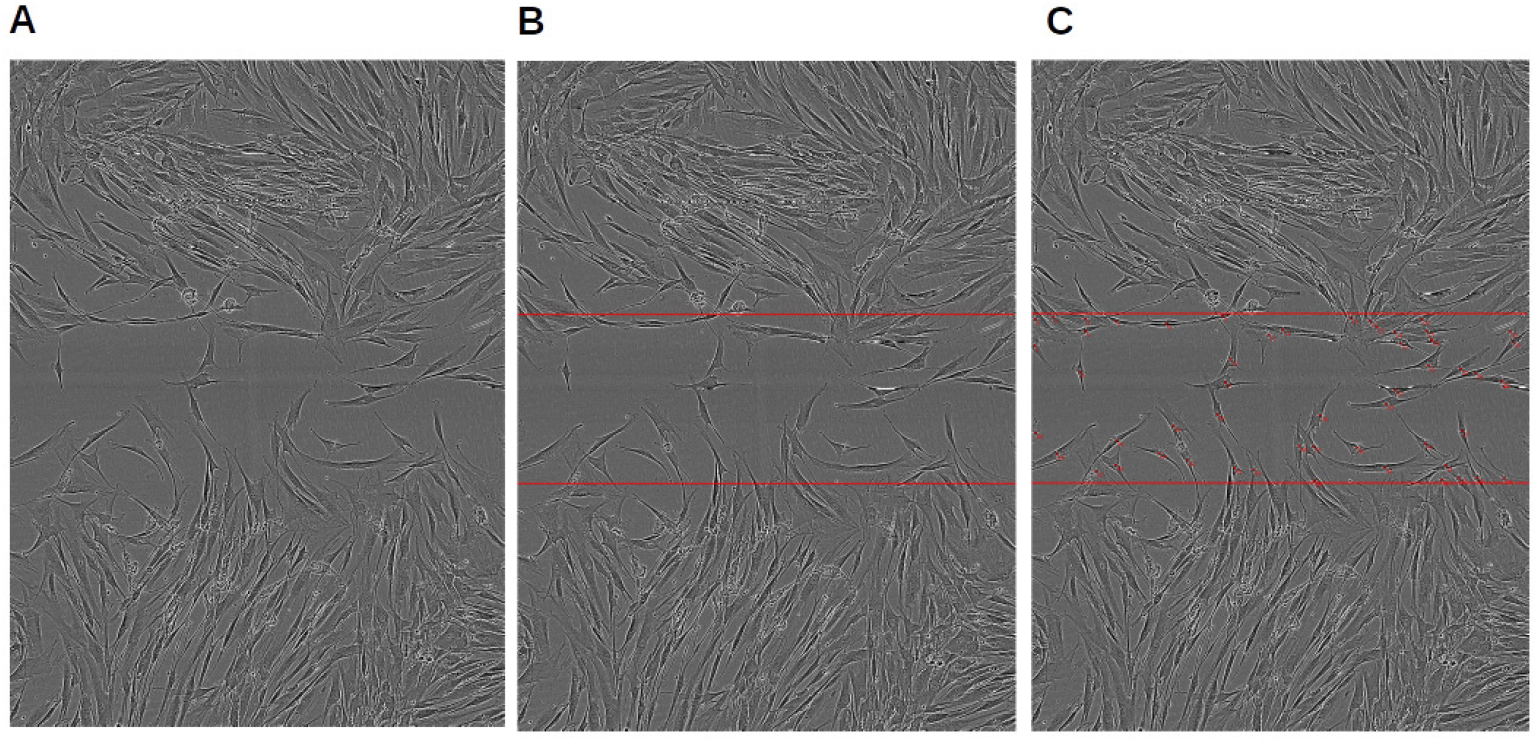
Experimental data quantification: CtrlA evaluation set. At *T*_0_ + 20 H, the experimental image is marked with the red lines define by (*x* ∈ [0; 1176] px, *y* = 600px) and (*x* ∈ [0; 1176] px, *y* = 1000px). This corresponds to the wound domain, empty at *T*_0_, to close by the fibroblasts. The fibroblasts inside this domain at *T*_0_ + 20 H, are labeled, contoured, and their surfaces measured. The sum of their surfaces (see Table 1) will be used as ground truth for the mathematical model calibration and evaluation.

In order to inform the mathematical modeling, the surface and the principal length of 231 control healthy fibroblasts and 245 keloid fibroblasts control group (CtrlA and B) are measured. These fibroblasts are selected by these following criteria: not overlapping with others fibroblasts, not located near the boundary of the picture, with a clear contrast of the cellular body. Fig.3 shows the resulting histograms on surface and Fig.4 on length. We note that keloid surfaces distribution is close to a Gamma distribution, with a ratio of 16 between the extreme values, whereas the healthy fibroblast surfaces distribution is less similar to a Gamma distribution, although the ratio between the extreme values is superior to 14. Regarding length distribution, keloid fibroblasts are closer to Gaussian distribution, and healthy fibroblasts more so (a ratio of 3 between the extreme values for healthy ones and 5 for keloids ones).

**Fig 3.**
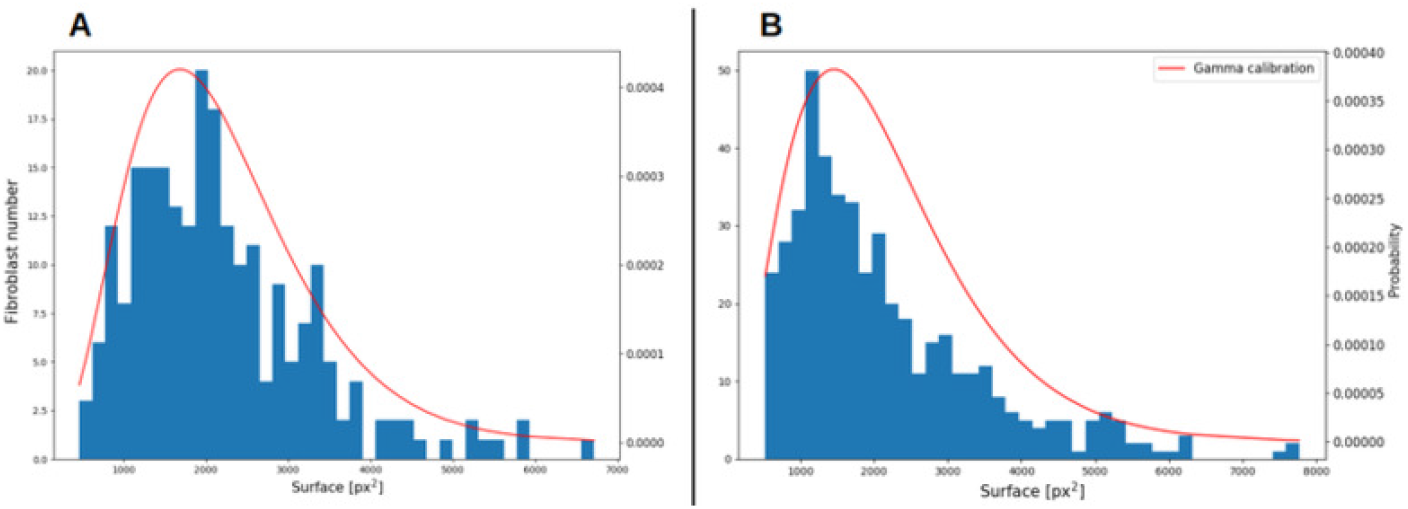
Experimental data quantification: surface measurements. **A** healthy fibroblasts; **B** keloid fibroblasts. Blue histogram: surface measurements of 231 control healthy fibroblasts and 471 keloid fibroblasts. Red line: we assume a Gamma distribution (see Eq.11) for the surface of both fibroblast types. The Gamma fit from SciPy library [30] gives *α* = 4.314 , *β* = 508.8 for healthy fibroblasts and *α* = 3.101 , *β* = 692.1 for keloid fibroblasts.

**Table 1.** Experimental data quantification: initial condition and progression of the wound closure.

| Assays | Nb. of fibroblast | % of wound closure at 20h | % of wound closure at 36h |
| --- | --- | --- | --- |
| HF-CtrlB | 352 | 18.29 | 31.90 |
| HF-CtrlA | 427 | 12.49 | 40.12 |
| CtrlA | 372 | 39.88 | 55.90 |
| CtrlB | 346 | 36.10 | 41.56 |
| M1A | 345 | 21.86 | 34.63 |
| M1B | 341 | 21.24 | 30.45 |
| M2A | 420 | 43.93 | 56.90 |
| M2B | 398 | 45.81 | 51.87 |

Column 2 (*T*_0_), all the fibroblasts’ nuclei are labeled for each experimental tests. Column 3-4 (*T*_0_ + 20 H, *T*_0_ + 36 H) We define, in the experimental images, a domain between the lines define by (*x* ∈ [0; 1176] px, *y* = 600px) and (*x* ∈ [0; 1176] px, *y* = 1000px) (see red lines Fig.2B,C), always empty at *T*_0_. This corresponds to the wound domain to close by the fibroblasts. The fibroblasts inside this domain at *T*_0_ + 20 H and *T*_0_ + 36 H are labeled, contoured, and their surfaces measured. The sum of the surfaces are provided in percentage of wound closure for each experimental tests. Compared to 6 keloid experiments, the healthy fibroblasts experiments shows a slower wound closure at *T*_0_ + 20 H and *T*_0_ + 36 H. Within the 6 keloids experiments, the M1 treatment largely slows down the wound closure (experiments M1A and B), and the M2 treatment slightly speeds it up at *T*_0_ + 20 H, but without significance at *T*_0_ + 36 H (experiments M2A and B).

### Agent-based modeling

#### Cellular Potts Model

Cellular Potts Model (CPM) was developed by Graner and Glazier in [11], and further extended in [31, 32] and many others. CPM allows a precise description of each cell geometry and the dynamics of the system is prescribed through an effective energy expression *ℋ*. Beforehand, this effective energy expression was termed ‘Hamiltonian’, and was changed in order to avoid confusion with balance laws of physics [12]. *ℋ* sums all the constrains applied the system, and its components depend on the targeted biological phenomenon.

Regarding scratch assays of human dermal fibroblasts without proliferation (blocked by mitomycin), we choose to focus on:

- active migration, *i.e*. haptotaxis, of the agent
- geometry constrains, length and surface, on the agent
- adhesive-repulsive contact based on agent to agent interaction

#### Constitutive equations

The contact constitutive law was designed to take into account the complex nature of fibroblast to fibroblast contact [23]. The contact law is first adhesive, since a small contact surface between two fibroblasts is shared. As the contact surface increase, the contact law is less adhesive and becomes repulsive since a threshold is reached. Follows the equation of the contact law *J*_c_ between fibroblasts *i* and the set of its neighbors *j*:

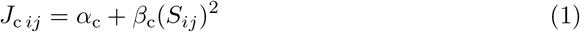

where *S*_*ij*_ is the shared surface between the fibroblast *i* and its neighbors *j, α*_c_ and *β*_c_ are constitutive parameters, left to be calibrated. Follows the corresponding term *J*_*c*_ of the effective energy expression:

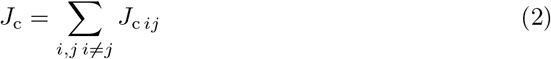

Following the same reasoning, the parameter *s*_targ *i*_, the targeted surface occupied of a fibroblast *i*, depends on the surface shared with its neighbors *S*_*ij*_ . This acknowledges the experimental observation that denser are the fibroblasts, smaller is the occupied surface by each individual.

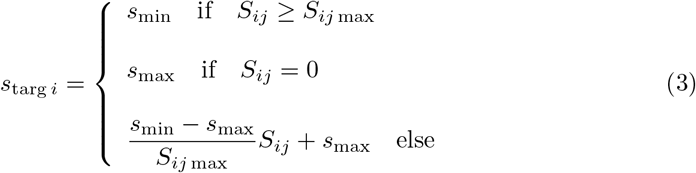

*s*_min_ = 600 px^2^ and *s*_max_ = 4800 px^2^ are deduced from experimental measurements. *S*_*ij* max_, left to be calibrated, is a constitutive parameter corresponding corresponding to an overcrowded configuration where the agent behavior has its expression inhibited. The corresponding term *J*_s_ of the effective energy expression is expressed as follow:

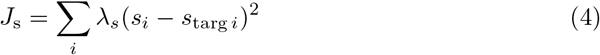

where *λ*_*s*_ is the weight of the surface constraint, left to be calibrated.

The elongation constrain *l*_targ *i*_ is built with the same function as *s*_targ *i*_, with *l*_min_ = 90 and *l*_max_ = 160 deduced from experimental data.

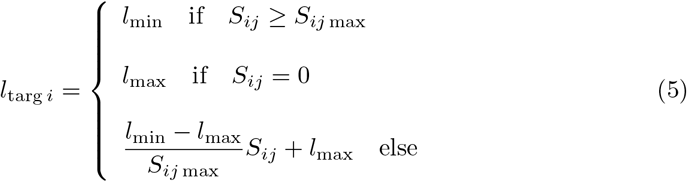

Follows the corresponding term *J*_l_ of the effective energy expression:

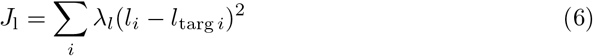

where *λ*_*l*_ is the weight of the elongation constraint, left to be calibrated.

#### Motility constitutive relationship: CIL hypothesis

Regarding agent motility, we test two constrains on healthy fibroblast and keloid fibroblast, to quantity the effect of the contact inhibition locomotion (CIL). If a fibroblast *i* is subjected to CIL, it obeys to the following form of motility constrain: *J*_m *i*_, defined by an isotropic displacement field **v**_**m**_, *i.e. v*_*m*_(*x*) = *v*_*m*_(*y*), prescribed to the center of mass of each agent, and subjected to the same function than *s*_targ_ and *l*_targ_:

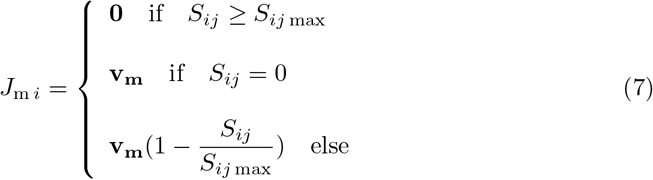

If we hypothesize that the motility of a fibroblast *i* is not sensitive to the contacts with its neighbors, *i.e*. not sensitive to CIL, the constrain is only defined by the isotropic displacement field **v**_**m**_:

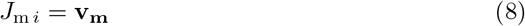

where the intensity of the displacement vector **v**_**m**_ is left to be calibrated. The motility constrain *J*_m_ then reads:

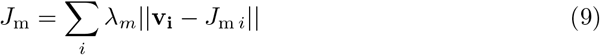

where the weight of the mobility constrain *λ*_*m*_ is left to be calibrated.

In the result section, we show the influence of the CIL on wound closures by keloid fibroblasts.

Summarize all the presented constrains, the effective energy expression *ℋ* corresponds to the sum of contact, surface, elongation and motility constraints:

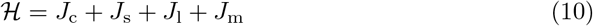

### Computational initial conditions

The pre-processed experimental data at *T*_0_ are used to design the numerical initial conditions. A lattice of the same resolution than the experimental image is created. The status of each pixel is informed by the following process:

- The nearest fibroblast nuclei label is found, and the corresponding polygon is provided (cf. to Fig.1A, B, C).
- The algorithm tests if the pixel is inside or outside the polygon. If yes, the pixel is labeled as type Fibroblast in the CPM, with the agent ID corresponding to the nuclei label.
- If no, the second nearest fibroblast nuclei label is tested. This process iterates up the third nearest nuclei, as further nuclei testing shows no improvement. If the tested pixel does not belong to any corresponding polygons, it is labeled as type Medium in the CPM, without an agent ID.

Fig.5 shows the resulting initial condition in the computational domain.

**Fig 4.**
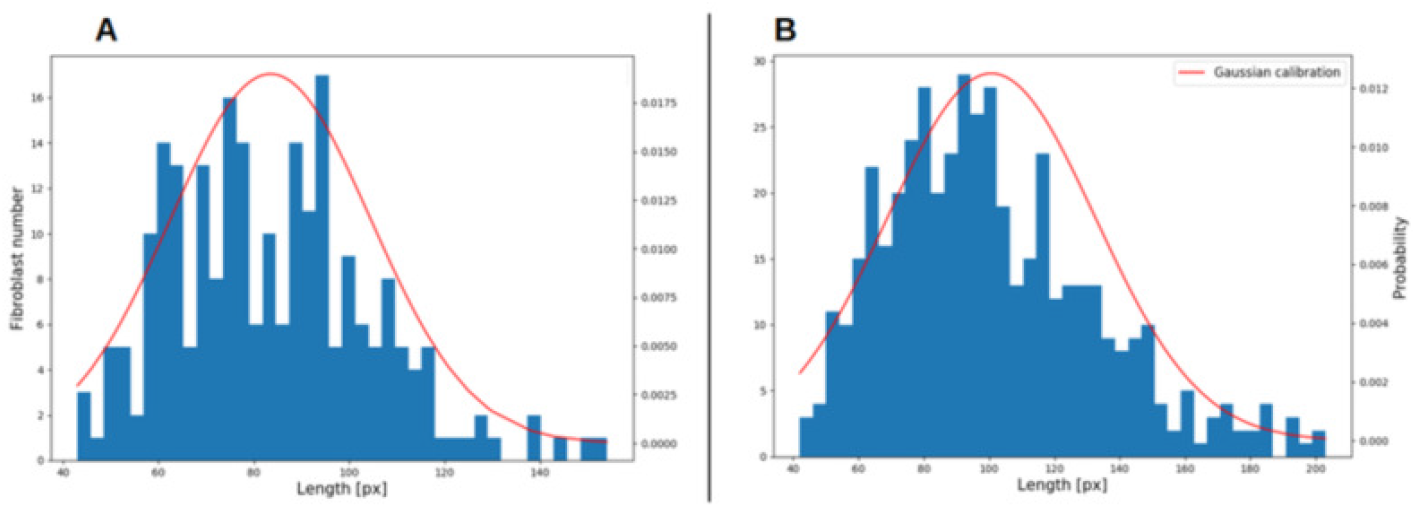
Experimental data quantification: length measurements. **A** healthy fibroblasts; **B** keloid fibroblasts. Blue histogram: length measurements of 231 control healthy fibroblasts and 471 keloid fibroblasts. Red line: we assume a Gaussian distribution for the length of both fibroblast types. The Normal fit from SciPy library [30] gives *μ* = 83.49 px , *σ* = 21.01 px for healthy fibroblasts and *μ* = 100.6 px , *σ* = 31.89 px for keloid fibroblasts.

**Fig 5.**
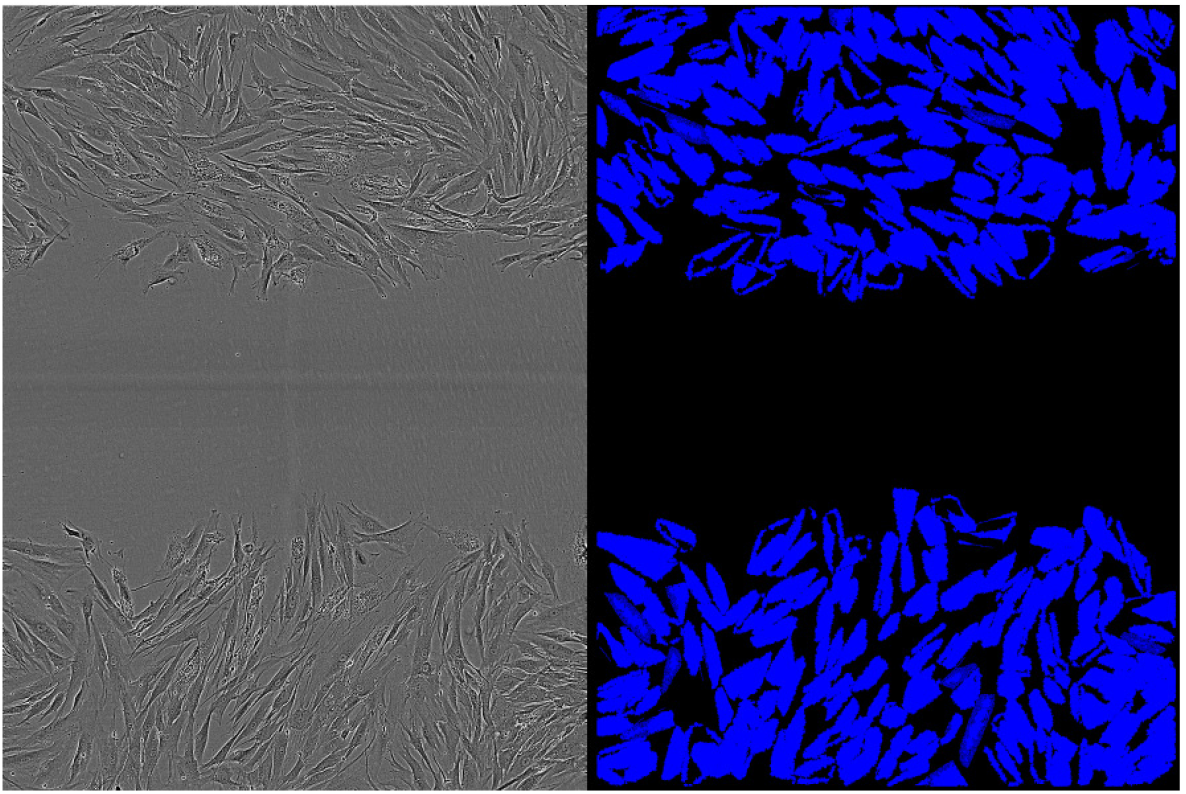
Numerical initial conditions from experimental image at *T*_0_. For each pixel, the nearest fibroblast nuclei label is found, and the corresponding polygon is provided (cf. to Fig.1A, B, C). If the pixel is inside the polygon, it is labeled as type Fibroblast in the CPM, with the agent ID corresponding to the nuclei label. If no, the second nearest fibroblast nuclei label is tested (up to the third), and the pixel is labelled the same way. If the tested pixel does not belong to any corresponding polygons, it is labeled as type Medium in the CPM, without an agent ID.

**Fig 6.**
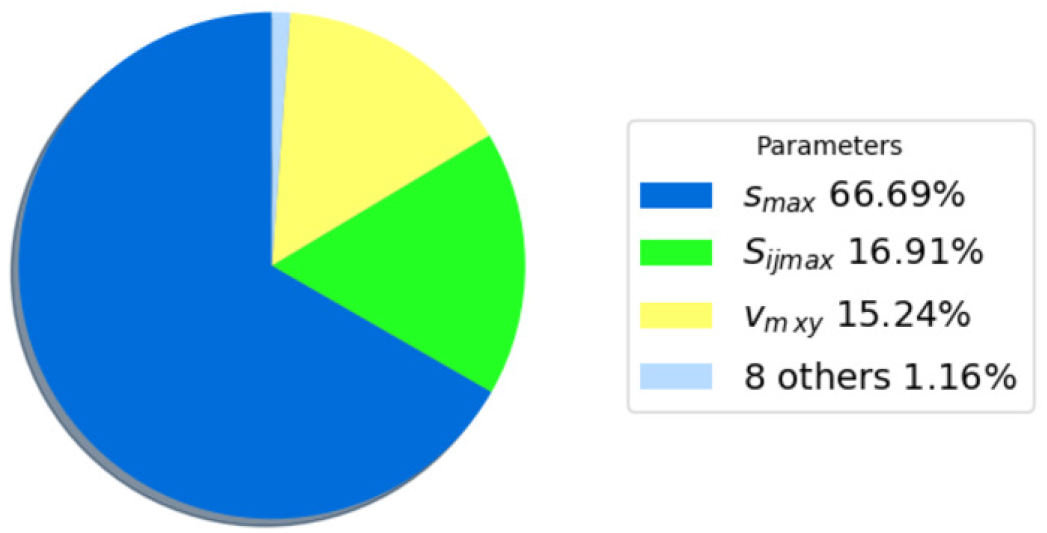
Sobol index of wound closure sensitivity on parameters. Only three parameters gather 98.84% of the solution variance: the parameters *s*_max_ (66.69%) the maximum surface that a fibroblast can occupy, *S*_*ij* max_ (16.91%) the maximum surface a fibroblast can shared with the set of its neighbors, and *v*_*m x*,*y*_ (15.24%) the isotropic motility of a fibroblast.

In order to provide initial values of targeted surface for each agent, the experimental measurements presented Fig.3 are fitted by a gamma distribution using SciPy library [30]. Therefore, the targeted surface *s*_targ *i*_ of each fibroblast *i* is initialized at the first time increment, following the Gamma distribution:

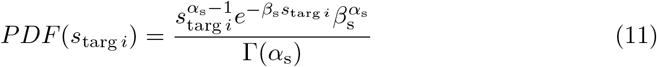

where Γ is the Gamma function, and *α, β* are obtained from the SciPy fit *α* = 4.314 , *β* = 508.8 for healthy fibroblasts and *α* = 3.101 , *β* = 692.1 for keloid fibroblasts.

### Computational framework

CPM simulations are run using CompuCell3D [12] version 4.0, with a single core of an Intel i5 8thGen. The code uses the following plugins: NeighborTracker, Contact, VolumeLocalFlex, PixelTracker, Motility and CenterOfMass presented in Swat *et al*. in [33], LengthConstraint and ConnectivityGlobal presented in Merks *et al*. in [31]. NeighborTracker plugin is used to build *S*_*ij*_, the shared surface between a fibroblast and the set of its neighbors. Contact plugin is used to define the constraint *J*_*c*_ Eqs.1,2. VolumeLocalFlex plugin is used to define the constraint *J*_*s*_ Eqs.3,4 with the *S*_*ij*_ dependency. PixelTracker plugin is used to make screenshots during the simulation. Motility plugin, and its dependency CenterOfMass plugin, are used to define the constraint *J*_*m*_ Eq.7 for healthy fibroblasts and Eq.8 for keloid fibroblasts.

LengthConstraint plugin, and its dependency ConnectivityGlobal, are used to build the constraint *J*_*l*_ Eqs.5,6. Only Contact, VolumeLocalFlex and LengthConstraint plugins were modified to correspond to the constitutive relationships Eqs.1, 3 and 5 presented in this study. These modifications are presented in Appendix and the full code is available at https://github.com/SUrcun/ABM_fibroblast.

## Results

Due to the initialization of the parameter *s*_targ_ with the Gamma distribution Eq.11, every computation was done three times. These three runs lead to a small results variation close but smaller than 1%. Therefore all the following results should be read with an uncertainty of *±*1%. The two first subsections *Influence of the CIL hypothesis on keloid fibroblasts* and *Local sensitivity analysis* are performed with the following initial set of 11 parameters:

*v*_*m x*,*y*_ = 5000; *λ*_*m*_ = 100; *S*_*ij* max_ = 130; *s*_min_ = 600; *s*_max_ = 4800; *λ*_*s*_ = 5; *l*_min_ = 50; *l*_max_ = 250; *λ*_*l*_ = 16; *α*_c_ = 100; *β*_c_ = 2.

After the sensitivity analysis, the parameter set is calibrated and evaluated on external data.

### Influence of the CIL hypothesis on keloid fibroblasts

The Table 2 shows wound closure percentage in the simulations of keloid fibroblasts with and without the CIL hypothesis. The numerical results of keloid fibroblasts without CIL are all closer to experiments than the ones with CIL, all parameters being equal. The overall dynamics of the simulations without CIL is closer to the experiments. First, the numerical results are less dependent on the total population of keloid fibroblasts. Second, the rate of wound closure per hour follows the same trend than the experiments: CtrlB shows 1.8% the first 20 hours, followed by 0.34% the next 16 hours; the simulation without CIL shows 1.31% and 0.23% respectively, whereas the simulation with CIL shows 0.93% and a decrease to 0.14%. The same trend is observable for the M1B and M2B simulations. Therefore, the sustained progression of keloid fibroblasts in wound healing may be simulated by the loss of CIL.

**Table 2.** Influence of the CIL hypothesis on keloid fibroblasts with the initial parameter set.

| Assays (Nb. of fibroblasts) | Time | Experimental | CIL | No CIL |
| --- | --- | --- | --- | --- |
| CtrlB<br>(352) | 20H | 36.10 | 18.60 | 26.18 |
|  | 36H | 41.56 | 20.90 | 29.99 |
| M1B<br>(341) | 20H | 21.24 | 19.27 | 25.29 |
|  | 36H | 30.45 | 22.40 | 36.43 |
| M2B<br>(398) | 20H | 45.81 | 32.05 | 34.75 |
|  | 36H | 51.87 | 34.24 | 44.78 |

### Local sensitivity analysis

We perform a local variance-based sensitivity analysis by Sobol index. For the details of this process, the reader is referred to Urcun *et al*. in [34]. Local sensitivity analysis is performed on CrtlB, M1B and M2B experiments. Each of the 11 parameters is perturbed one at a time starting from its initial value. The resulting Sobol indices show that 3 parameters gather 98.84% of the solution variance, with the parameter *s*_max_ dominant (66.69% of the variance), see Fig.6. The 8 other parameters are therefore considered fixed and the 3 governing parameters, *s*_max_, *S*_*ij* max_ and *v*_*m x*,*y*_ will be calibrated. For the details of the Sobol indices, see in Appendix (Table 5).

### Calibration

To measure the distance between experimental data and numerical results, we followed the prescription of [35]: the root mean square error (RMSE) relative to a reference, *i.e*. normalized RMSE (nRMSE). The nRMSE of the numerical closure C_num_ relative to the experimental closure C_data_ is computed as:

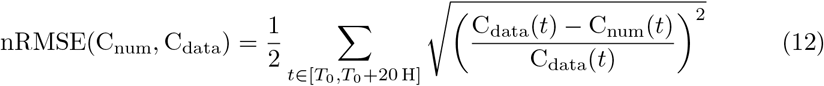

The three governing parameters *s*_max_, *S*_*ij* max_ and *v*_*m x*,*y*_ are calibrated by user because the cost function knows multiple local minima, gradient-based optimization methods are, as far as we know, not available in CC3D and global gradient-free methods, such as evolutionary or genetic algorithms were prohibited due to computational cost: one iteration until *T*_0_ + 36 H takes 10 hours of computational time in the current configuration. Therefore, if the calibration of the control experiments HF-CtrlB (41 iterations) and CtrlB (78 iterations) show a good agreement with experimental data at 5.60% and 4.02% nRMSE respectively, we cannot ensure it is their global minimum. After calibration of CtrlB experiment, a M1 experiment was calibrated (8 iterations) with the priority of a decreased motility, as described by experimental findings. In parallel, a M2 experiment was calibrated (21 iterations). Quantitative results of the calibration are shown Table 3, and are illustrated Fig.7. Calibration of M1 treatment leads to a 8.4% decrease of agent maximum surface *s*_max_, a 6.6% decrease of the agent *i* maximum shared surface with its *j* neighbors *S*_*ij* max_ and a 33% decrease of the agent motility *v*_*m x*,*y*_. Excellent calibration of M2 treatment (0.45% nRMSE) is obtained only by a 0.7% increase of agent maximum surface *s*_max_.

**Fig 7.**
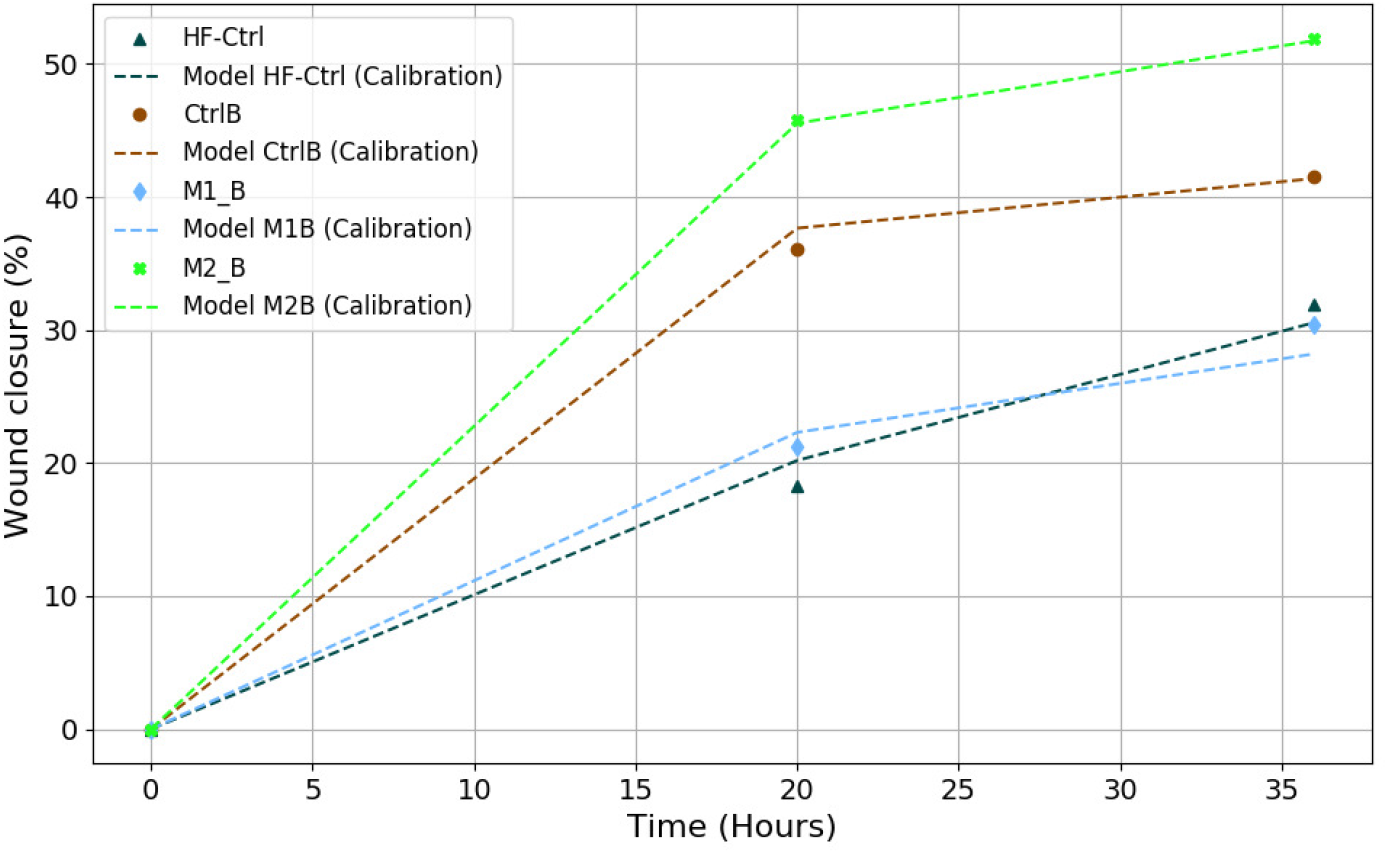
Experimental data against calibrated model parameters. Numerical results of CtrlB (brown, dotted) shows a good agreement with experimental data (brown, circle) 4.02% nRMSE; numerical results of M1 treatment M1B (light blue, dotted), obtained by decreased motility and decreased shared surface of CtrlB parameters, shows a good agreement with experimental data (light blue, diamond) 6.30% nRMSE; numerical results of M2 treatment M2B (light green, dotted), obtained by a slight increased motility of the CtrlB parameters, shows a excellent agreement with experimental data (light green, square) 0.45% nRMSE.

**Table 3.**
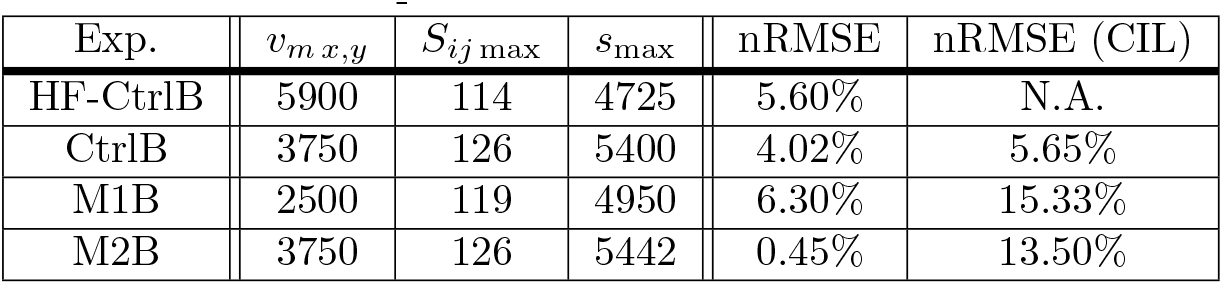
Calibrated sets of parameters. The three governing parameters, *s*_max_, *S*_*ij* max_ and *v*_*m x*,*y*_, are calibrated for CtrlB experiment, then a M1 experiment and a M2 experiment are calibrated in parallel. The M1 calibration provokes a 33% decrease of the agent motility *v*_*m x*,*y*_. The excellent M2 calibration is obtained only by a 0.7% increase of agent maximum surface *s*_max_.

### Evaluation

The 3 calibrated parameters are evaluated against the second half of the experimental dataset, namely HF-CtrlA, CtrlA, M1A and M2A. With these new initial conditions, we show that the mathematical modeling is compliant to reproduce the same experimental settings, with the inherent variability of *in vitro* experiments. Although the model with HF-CtrlA conditions shows the largest error, 18.75% nRMSE, with experimental data, both experimental and numerical results follow the same trend: the increased population of HF-CtrlA (427 fibroblasts *vs*. 352 in calibration) provokes an overcrowd phenomenon that drops off the healing progression at *T*_0_ + 20 H, before accelerates. The evaluated dataset shows a good agreement with the keloid experimental data, see Fig.8. The error on the control group CtrlA shows an increase of 6.81%, and the error on the M2 secretome group M2A increases of 8.33%. We note that the error on the M1 secretome group M1A shows a larger increase with 10.54%, see Table 4.

**Fig 8.**
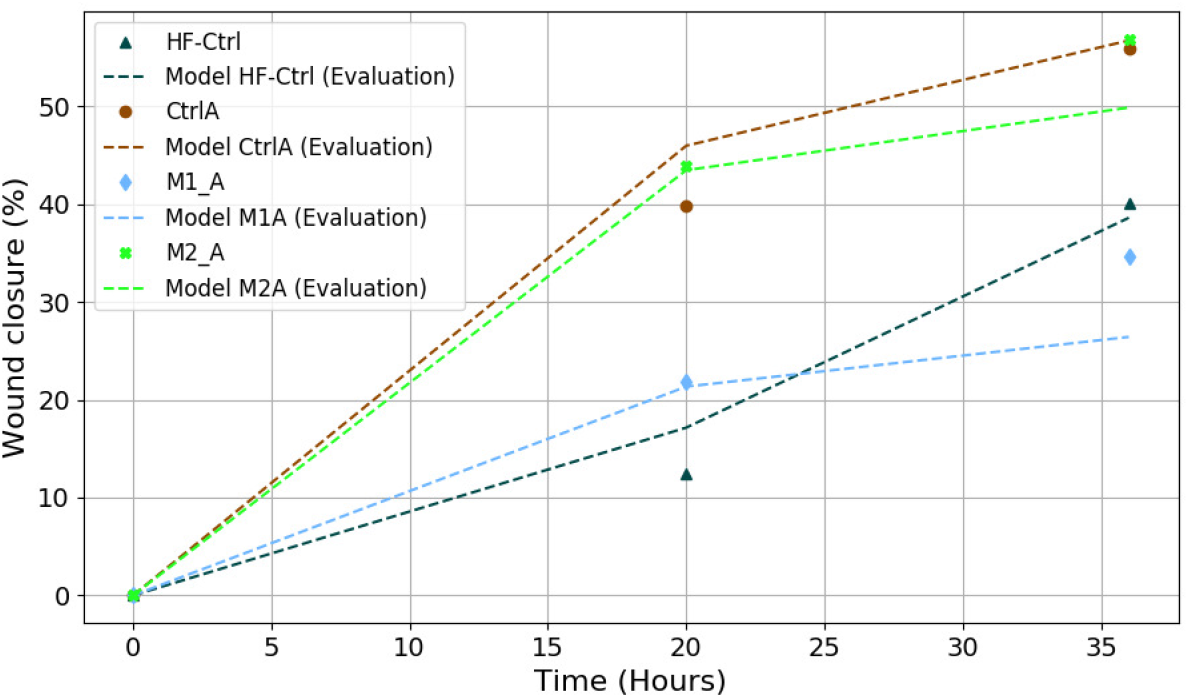
Experimental data against evaluated model parameters. Evaluation of control parameters on HF-CtrlA (black, dotted) shows the largest error, 18.75% nRMSE, with experimental data (black, triangle). However, the model shows the same overcrowd phenomenon - dropped off at *T*_0_ + 20 H, then acceleration - of the experimental data. Evaluation of control parameters on CtrlA (brown, dotted) shows a reasonable agreement with experimental data (brown, circle) 4.02% nRMSE; Evaluation of M1 treatment parameters M1A (light blue, dotted) shows an error with experimental data (light blue, diamond) of 16.84% nRMSE; Evaluation of M2 treatment parameters M2A (light green, dotted) shows a reasonable agreement with experimental data (light green, square) 8.78% nRMSE.

**Table 4.** Evaluation of the calibrated sets of parameters.

| Exp. | $v_{m\ x,y}$ | $S_{ij\ \max}$ | $s_{\max}$ | nRMSE | nRMSE (CIL) |
| --- | --- | --- | --- | --- | --- |
| HF-CtrlA | 5900 | 114 | 4725 | 18.75% | N.A. |
| CtrlA | 3750 | 126 | 5400 | 10.83% | 8.04% |
| M1A | 2500 | 119 | 4950 | 16.84% | 21.06% |
| M2A | 3750 | 126 | 5442 | 8.78% | 19.24% |

## Discussion

In the research field of wound healing, scratch assays are fundamental experiments. If we aim to a robust modeling of these experiments, it should contain: i) an reliable error quantification between experiments and numerical results, ii) a model that accurately reproduce the experiment, iii) with a number of parameters small enough to be quickly adapted to the inherent diversity of these experiments. With only 3 governing parameters (98.84% of the solution variance) and CIL for physiological behavior, our model is able to reproduce scratch assays of human dermal fibroblasts, healthy and keloid, and their variations induced by macrophage secretome treatment. The calibration gives a very good agreement with the experimental data with an error from 0.45% to 6.30 *±* 1% nRMSE. The evaluation dataset shows a reasonable agreement with the experimental data with an increase of error from 8.87% to 16.84 *±* 1% nRMSE for keloid fibroblast and a larger error of 18.75 *±* 1% nRMSE for healthy fibroblast.

The calibrated parameter *s*_max_ - the maximum surface that a fibroblast can occupy - is in accordance the experimental measurements, as its value corresponds to the upper part these measurements. The motility parameter *v*_*m x*,*y*_ is calibrated according to the experimental findings: M1 secretome treatment decreases the fibroblasts’ motility. The only governing parameter that remains phenomenological is *S*_*ij* max_ - the maximum shared surface between a fibroblast and the set of its neighbors - because, to our knowledge, there is a gap of experimental findings on fibroblast to fibroblast contact dynamics. The only study we found, detailed but only qualitative, was from Chen *et al*. [23] in 1982. New experimental studies that can quantify this dynamics, coined as contact inhibition of locomotion (CIL) by Abercombie *et al*. in 1953 [22], would be highly valuable. In the agent-based model, the only difference between healthy and keloid fibroblasts is the motility behavior. Following the claim of Mayor *et al*. in [24], malignant cells lose their sensitivity to CIL, therefore the motility of keloid fibroblast is not dependent on *S*_*ij* max_, *i.e*. not dependent on the shared surface with its neighbors. In the numerical results, we show that wound closure of healthy fibroblast follows almost a linear trend in HF-CtrlB (see Fig.7). When fibroblasts density increases in HF-CtrlA (see Fig.8), the wound closure slows down before accelerating. This could be interpreted as an overcrowd phenomenon [25], and reinforce the hypothesis of CIL in healthy fibroblast. Even with a larger error, this phenomenon is reproduced by our model. On the other hand, the 6 experiments of keloids fibroblasts do not show the phenomenon, whatever their population density. To test our hypothesis, we show the numerical results of keloid subjected to CIL. These results always display a greater error compared to experiments, both with the initial parameter set extracted from experiments and the calibrated set. Only one exception occurs in the external evaluation dataset (8.04% *vs*. 10.83% for the keloid control experiment).

One drawback of the current implementation is the manual assimilation of experimental data. This time-consuming task impedes to reproduce these results on a large experimental dataset. To alleviate this problem, some collaborators are currently working on a tailored machine learning application for fibroblast counting and body contouring. The second drawback is the computational cost due to the single-core configuration: 10 hours of computational time for 36 hours of simulated time. This configuration impedes a global optimization, and this current situation may be solved by an efficient parallel implementation of the CCD3 solver.

In future studies, this agent-based model will be used as input parameter for a upscale model, currently under development, to informed a physical model at the tissue scale.

## Acknowledgments

SPAB and SU acknowledge funding from the Luxembourg National Research Fund (FNR) grant number INTER/ANR/21/16399490; GR acknowledges funding from the ANR grant number ANR-21-CE45-0025-03; RE and AL acknowledge funding from the French Agence Nationale de la Recherche (ANR) grant number ANR-21-CE45-0025-01.

### Appendix

#### Examples of CC3D codes

Modification of the Contact plugin

~~~
    <Plugin Name=“ Contact “>
                 <Energy Type1=“Medium” Type2=“Medium”>0</ Energy>
                 <Energy Type1=“ Fibro blast “ Type2=“ F i b r o b l a s t “
                 id=“ neighbor dependent “>100</ Energy>
                 <Energy Type1=“ Fibro blast “ Type2=“Medium”>20</ Energy>
                 <Neighbor Order>3</ Neighbor Order>
    </ Plugin>
~~~

The fibroblast-fibroblast contact law is modified according to Eq.1 through the variable neighbor dependent, implemented in the following Python code:

~~~
**class** Neighbor Tracker Printer Steppable ( Steppable Base Py ) :
   **def __**init__ (self, f requency =1 ):
      Steppable Base Py . __init__ (self , f requency )
   **def** step (self , mcs ) :
      **for** cell **in** self . cell_list :
        neighbour_list = self . get_cell_neighbor_data_list (cell)
        neighbour_dependent = self . get_xml_e l ement (
        ‘ neighbour_dependent ‘ )
      **for** neighbor , common_surface_area **in** neighbor_list :
         **if** neighbor :
           neighbour_dependent . cdata=−a lpha+beta *
           **float** (neighbour_list . commonSurfaceAreaWithCellTypes (
           cell_type_list =[ 1 , 2 ] ) ) *
           **float** (neighbour_lis t . commonSurfaceAreaWithCellTypes (
           cell_type_list =[ 1 , 2 ] ) )
~~~

Modification of the VolumeLocalFlex plugin. The influence of fibroblast-fibroblast contact surface on target surface for each agent is implemented according to Eq.3, in the following Python code:

~~~
**class** VolumeParamSteppable ( Steppable Base Py ) :
   **def __**init__ ( self , frequency =1 ):
      Steppable Base Py . ___init__ (self , frequency )
   **def** start (self) :
     **for** cell **in** self . cell list :
       **if** (cell . **type**==1):
         initial_volume = **int** ( np . random . gamma( 3 . 7 , scale =1370 ,
         size =1))
         cell . target Volume = initial volume
         cell . lambdaVolume = 5 . 0
   **def** s tep ( self , mcs ) :
     **for** cell **in** self . cell list :
       **if** (cell . **type**==1):
         neighbour_li s t = self . get_cell_neighbor_data_list ( cell )
     **for** neighbor , common_surface_area **in** nei ghbor_list :
        **if** neighbor :
           Sij=**float** (neighbour_list . commonSurfaceAreaWithCellTypes (
           cell_type list =[ 1 , 2 ] ) )
          **if** Sij >maxshared :
             cell . target Volume = minsurf
          **elif** Sij ==0:
             cell . target Volume = maxsurf
          **else** :
            cell . target Volume = **int** ( **round** ( ( ((
            minsurf −maxsurf )/ maxshared ) * Sij+maxsurf ) ))
~~~

Modification of the LengthConstraint plugin. The influence of fibroblast-fibroblast contact surface on target length for each agent is implemented according to Eq.5, in the following Python code:

~~~
**class** Elongation Flex Steppable ( Steppable Base Py ) :
   **def __**init__ (self , frequency =1 ):
      Steppable Base Py . __init__ (self , frequency )
   **def** start (self ) :
     **for** cell **in** self . cell_list :
       **if** cell . **type** == 1 :
           target Length= 160
           self . length Constraint Plugi n . set Length Constraint Data (
           cell , 16 , target Length )
           cell . connectivity On = True
   **def** step (self , mcs ) :
     **for** cell **in** self . cell list :
       **if** (cell . **type**==1):
          neighbor list = self . get_cell_neighbor_data_list (cell )
   **for** neighbor , common surface area **in** neighbor list :
     **if** neighbor :
       Sij=**float** (neighbour_list . commonSurfaceAreaWithCellTypes (
       cell type list =[ 1 , 2 ] ) )
      **if** Sij >maxshared :
        self . le ngth Constraint Plugin . set Length Constraint Data (
        cell , 1 6 , minlength )
      **elif** Sij ==0:
        self . length Constraint Plugin . set Length Constraint Data (
        cell , 1 6 , maxlength )
      **else** :
        self . length Constraint Plugin . set Length Constraint Data (
        cell , 16 , **int** ( **round** ( ( (( minlength −maxlength )/ maxshared ) *
        Sij+maxlength ) ) ) )
~~~

#### Results tables of sensitivity analysis, calibration and evaluation

**Table 5.** Detailed results of first order wound closure sensitivity on parameters with a variation of *±*10 %.

| Parameter | $\theta$ | Sobol index [%] |
| --- | --- | --- |
| $S_{ij_{\max}}$ | 2.0945 | 15.24 |
| $\lambda_s$ | -0.475 | 0.78 |
| $\alpha_c$ | -0.2345 | 0.20 |
| $\beta_c$ | -0.1495 | 0.08 |
| $\lambda_m$ | -0.1031 | 0.03 |
| $l_{\max}$ | 0.03 | 0.003 |
| $v_{m\,x,y}$ | -2.206 | 16.91 |
| $s_{\max}$ | 4.381 | 66.69 |
| $l_{\min}$ | -0.0812 | 0.02 |
| $s_{\min}$ | -0.021 | 0.002 |
| $\lambda_l$ | 0.0935 | 0.03 |

**Table 6.** Calibration numerical results by keloid fibroblast with and without CIL hypothesis.

| Assays | Hypothesis | % at 20h | % at 36h |
| --- | --- | --- | --- |
| CtrlB | No CIL | 37.65 | 41.38 |
|  | CIL | 39.42 | 44.30 |
| M1B | No CIL | 22.30 | 28.20 |
|  | CIL | 20.29 | 21.21 |
| M2B | No CIL | 45.54 | 51.74 |
|  | CIL | 36.62 | 42.49 |

**Table 7.** Evaluation numerical results by keloid fibroblast with and without CIL hypothesis.

| Assays | Hypothesis | % at 20h | % at 36h |
| --- | --- | --- | --- |
| CtrlA | No CIL | 45.96 | 56.72 |
|  | CIL | 46.14 | 57.90 |
| M1A | No CIL | 21.34 | 26.42 |
|  | CIL | 20.26 | 21.84 |
| M2A | No CIL | 43.46 | 49.86 |
|  | CIL | 39.79 | 45.84 |

